# Grapevine rootstocks affect growth-related scion phenotypes

**DOI:** 10.1101/864850

**Authors:** Zoë Migicovsky, Peter Cousins, Lindsay M. Jordan, Sean Myles, Richard Keith Striegler, Paul Verdegaal, Daniel H. Chitwood

**Affiliations:** Department of Plant, Food and Environmental Sciences, Faculty of Agriculture, Dalhousie University, Truro, NS B2N 5E3, Canada; E. & J. Gallo Winery, Modesto, CA, 95354, USA; E. & J. Gallo Winery, Acampo, CA 95220, USA; University of California Cooperative Extension, San Joaquin Valley, Stockton, CA, 95206, USA; Department of Horticulture, Michigan State University, East Lansing, Michigan, 48824, USA; Department of Computational Mathematics, Science and Engineering, Michigan State University, East Lansing, Michigan, 48824, USA

**Keywords:** rootstocks, grafting, grapevines, root-shoot interactions

## Abstract

Grape growers use rootstocks to provide protection against pests and pathogens and to modulate viticulture performance such as shoot growth. Our study examined two grapevine scion varieties (‘Chardonnay’ and ‘Cabernet Sauvignon’) grafted to 15 different rootstocks and determined the effect of rootstocks on eight traits important to viticulture. We assessed the vines across five years and identified both year and variety as contributing strongly to trait variation. The effect of rootstock was relatively consistent across years and varieties, explaining between 8.99% and 9.78% of the variation in growth-related traits including yield, pruning weight, berry weight, and Ravaz index (yield to pruning weight ratio). Increases in yield due to rootstock were generally the result of increases in berry weight, likely due to increased water uptake by vines grafted to a particular rootstock. We demonstrated a greater than 50% increase in yield, pruning weight, or Ravaz index by choosing the optimal rootstock, indicating that rootstock choice is crucial for grape growers looking to improve vine performance.

## Introduction

Grafting joins two distinct plant parts: a scion (shoot system) from a donor plant and a rootstock (root system) from a second plant to which the scion is attached. The practice of grafting chiefly enables clonal propagation but can also have many other benefits, such as reducing the juvenility period (increasing precocity) or size (dwarfing) in fruit trees^1–3^.

In grapevines (*Vitis vinifera* L.), widespread use of grafting began in the late 1800s, following the introduction of phylloxera (*Daktulosphaira vitifoliae* Fitch) to Europe from North America. While *V. vinifera* is highly susceptible to phylloxera, which feeds on the roots of grapevines, eastern North American *Vitis* species evolved in the presence of phylloxera and are tolerant and/or resistant to it. By grafting *V. vinifera* scions to rootstocks of other *Vitis* species, *V. vinifera* could be grown in European soils containing phylloxera, rescuing the wine industry^4^.

Ten years after its detection in Europe, own-rooted (ungrafted) grapevines with phylloxera were first identified in California. The inter-continental spread of the pest was likely due to the importation of vines from European nurseries or from eastern North America^5^. However, due to the sandy soils of California’s Central Valley (or San Joaquin Valley, specifically), phylloxera infections were not as severe and did not require the immediate use of rootstocks^6^. By the 1950s, less than 30% of California grapevines were grafted onto phylloxera-resistant rootstocks^7^. Still, over time, the California grapevine industry transitioned primarily to grafted vines. Currently, more than 80% of vineyards worldwide grow grafted vines^4^.

In addition to allowing *V. vinifera* vines to grow in phylloxera-infested soils, grapevine rootstocks can provide tolerance to several other damaging pests and diseases including root-knot and dagger nematodes^8–10^. Rootstocks may also be used to improve resilience to abiotic stresses such as salinity^11^ and drought^12^. Grafting grapevines to a particular rootstock can influence a wide range of traits in the scion including mineral composition^13–15^, berry chemistry^16^, and berry maturation^17^.

Of particular interest to grape growers is the observation that rootstock choice can affect vine size and yield^18^. While other factors such as climate and location exceed the influence of rootstock on grapevine growth^19,20^, numerous studies have provided evidence of the impact rootstock can have on yield^18,21,22^. For a grape grower, an increase in yield is desirable, but increasing vine size or vegetative growth also increases the cost of managing the vine, due to additional labour for vine training, pruning, and fruit thinning. An ideal rootstock will increase reproductive growth, or yield, without an accompanying increase in vegetative growth, which is assessed by measuring pruning weight or the amount of one-year-old dormant cuttings removed during the winter. The Ravaz index, or yield divided by pruning weight from the following dormant season, can be calculated to determine the relative ratio of reproductive to vegetative growth. The impact that rootstocks can have on berry composition is generally thought to be an indirect effect as a result of their impact on vegetative and reproductive growth, for example by altering water or nutrient uptake^19,23^.

With all the potential benefits offered by a rootstock, deciding which one to use is an important choice. While other changes to vineyard management can be made throughout the lifespan of the vines–such as altering irrigation, fertilizer, pesticides, and pruning–rootstock choice is made only once. In this study, we assessed eight traits of viticultural importance across two scion varieties (‘Chardonnay’ and ‘Cabernet Sauvignon’) grafted to 15 different rootstocks. The vines were grafted near Lodi in San Joaquin County, California, in 1992 and evaluated from 1995 to 1999 in order to determine the relative contributions of variety, year, and rootstock to phenotypic variation.

## Materials and Methods

### Experimental design

In 1991, dormant field grown rootstocks were planted in a Tokay fine sandy loam soil^17^. On April 10th, 1992, scionwood was whip-grafted to the planted rootstock. Rows were oriented east-west with vine spacing of 2.13 m by 3.05 m (Figure S1). The trellis system was a bilateral cordon with fixed foliage wires. The cordon wire was at 1.07 m height with single foliage wire about 40.6 cm above. There were two wires 45.7 cm above the foliage catch wire at either ends of a 63.5 cm cross arm. The vines were cordon trained and spur pruned.

Prior to vineyard establishment, wine grapes were grown at the site for over 75 years. Initial plantings on this site were ungrafted *V. vinifera* vines. Because of this production history, various pests were considered to be endemic. These included several species of nematodes, phylloxera, many grape associated viruses, and oak root fungus (*Armillaria mellea)*^18^. All of these soil pests and pathogens can cause considerable economic losses to growers. For this reason, ungrafted vines were not included as a control in this study.

Vines were grafted to the following rootstocks: ‘Freedom’^24^, ‘Ramsey’, ‘1103 Paulsen’, ‘775 Paulsen’, ‘110 Richter’, ‘3309 Couderc’, ‘Kober 5BB’, ‘SO4’, ‘Teleki 5C’, ‘101-14 Mgt’, ‘039-16’^25^, ‘140 Ruggeri’, ‘Schwarzman’, ‘420 A’, and ‘K51-32’^26^. The two scion varieties were ‘Chardonnay’ (selection FPS 04) and ‘Cabernet Sauvignon’ (selection FPS 07).

The experimental design was a randomized complete block design, split between ‘Chardonnay’ and ‘Cabernet Sauvignon’. There were four replications per treatment (rootstock). There were eight or nine vines per plot, except for ‘Kober 5BB’ and ‘SO4’, which had four or five vines each, to fit all treatments in the block. Data were collected for five years from 1995-1999.

### Vine management

Canopy management practices were consistent with regional guidelines and included shoot thinning and leaf removal. Shoot thinning was performed pre-bloom and consisted of removal of non-count shoots (shoots not originating from spur positions). Leaf removal was performed at berry set on the north side of the vine only, to avoid excessive exposure and possible sunburn on the south side. Four to six leaves were removed to open a window in the fruiting zone.

Irrigation and vine nutrition programs were standard Best Management Practice for the Lodi District. The irrigation strategy followed a moderate Regulated Deficit Irrigation (RDI) program of about 80% estimated crop evapotranspiration (ETc) losses, from berry set to veraison^19^.

During the post-harvest period, vineyard irrigation was increased to 100% ETc. The vine nutrition program consisted of the application of approximately 30 lbs of actual nitrogen (N) and 60 lbs of actual potassium (K) per acre at post bloom annually. Zinc (Zn) was applied in some years, as local soils tend to be low in native levels of Zn^20^. All irrigation and nutrients were applied through a drip system, composed of two 0.5 gallons per hour emitters per vine.

### Data collection

Prior to harvest, a 100 berry sample was collected for each plot. The berries were counted and weighed to determine average berry weight. Berries were crushed by hand in plastic collection bags, then strained through cheesecloth to provide juice for analysis of soluble solids content (SSC) (°Brix), pH, and titratable acidity (TA) (g/L). Juice samples were titrated to an endpoint of pH 8.2 to determine TA^21^. SSC was determined by a temperature compensating Atago N1 refractometer (20 °C) and pH was measured using Beckman 200 pH meter with a dual KCl electrode. Grapes were harvested once they reached an acceptable commercial level for SSC, approximately 24.5 °Brix for ‘Cabernet Sauvignon’ and 23 °Brix for ‘Chardonnay’. Within a particular variety (‘Cabernet Sauvignon’ or ‘Chardonnay’) all vines were harvested on the same day (Table S1).

The number of clusters per vine and total fruit yield were recorded. In late winter, vines were pruned to retain two node fruiting spurs with a target of 50 nodes retained per vine (Table S1). Dormant pruning weights were measured.

Weather data from 1994 to 1999 were downloaded from the National Environmental Satellite, Data, and Information Service for Lodi, California, US (USC00045032) on September 30, 2019. Minimum temperature, maximum temperature, and cumulative precipitation for 1994 to 1999 were plotted (Figure S2).

### Statistical analysis

We calculated Ravaz index, a measurement of crop load, by dividing yield by pruning weight from the following dormant season. As a result, our dataset consisted of eight traits, measured across five years, for two scion varieties grafted to 15 different rootstocks (Table S2). The experimental design included a replication term (block) to indicate the position of the vines in the vineyard, which is a randomized complete block design, as evidenced by Figure S1.

The following linear model (Equation 1) was evaluated for each phenotype:

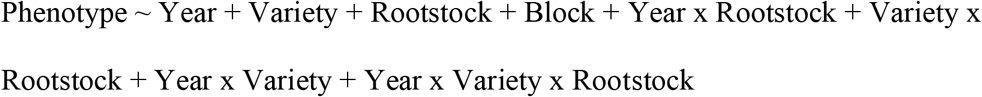

The model was optimized for each phenotype, which included the removal of the three-way interaction in all cases as well as non-significant two-way interactions. All main effects were retained. The code used for analyzing and visualizing the data in this study can be found on GitHub^27^. All terms in the model were fixed and the analysis was performed in R using the lm() and aov() functions from the stats package^28^. After running the model, the distribution of the residuals was examined to check for normality. Next, data were tidied using the tidy() function from the broom R package^29^. The percent variation was calculated for all terms by calculating the Sum of Squares for a particular term, divided by the Total Sum of Squares, then multiplied by 100. The results for significant terms (p < 0.05), except block (position in the vineyard), were plotted. We included block in our model to account for variation due to position in the vineyard, but we do not discuss those results here. They are included in our supplemental files and explain up to 10.72% of the variation in a trait (Table S3).

We visualized phenotype data for ‘Chardonnay’ and ‘Cabernet Sauvignon’ separately using a loess smoothing line to plot variation across years. For the four traits where rootstock explained the largest amount of variation (i.e. yield, berry weight, pruning weight, and Ravaz index), rootstocks were compared using a Tukey Test on the model results. For each phenotype, the raw data and a corresponding boxplot were plotted for each rootstock. The estimated marginal means and corresponding 95% confidence intervals were calculated and plotted using the emmeans package version 1.5.1 in R^30^ To visualize the variety-specific rootstock effects, we plotted the median values (+/- standard deviation) for ‘Cabernet Sauvignon’ and ‘Chardonnay’ separately for each phenotype (Figure S4).

Since there are large differences between the two grape varieties used in this study, we calculated the correlation between phenotypes for ‘Chardonnay’ and ‘Cabernet Sauvignon’ separately, using a Spearman’s correlation in R v.3.60^22^. To correct for multiple testing, p-values within a particular variety were Bonferroni-corrected. Heatmaps were generated using the heatmap.2 function in the gplots R package^23^. All remaining figures were plotted using ggplot2 in R^24^.

Lastly, we determined the potential range of variation induced by rootstock choice by calculating the percent change possible from the lowest rootstock median value to the highest rootstock median value within a particular phenotype. These results were visualized with phenotypes ordered from highest to lowest possible percent change.

## Results

### Phenotype variation across years

There was strong variation in phenotypes across the years of the study (Figure 1, Figure S5). The average yield across all rootstocks decreased for both varieties in 1996 (average of 11.3 Kg for ‘Cabernet Sauvignon’ and 7.9 Kg for ‘Chardonnay’) and 1998 (average of 8.53 Kg for ‘Cabernet Sauvignon’ and 8.6 Kg for ‘Chardonnay’) in contrast to other years where yields ranged from 10.4 Kg to 15.3 Kg for ‘Chardonnay’ and 13.3 Kg to 17.3 Kg for ‘Cabernet Sauvignon’, with the highest yields for both varieties produced in 1997. Similarly, the average number of clusters for either variety was lowest in 1996 and 1998 for ‘Cabernet Sauvignon’ with values of 80.4 and 71.8, respectively, in contrast to other years where values ranged from 111 to 133. For ‘Chardonnay’, the lowest number of clusters, on average, was produced in 1998 (59.9) and although many rootstocks had lower numbers in 1996, the overall average was slightly higher (69.5) than 1995 (63.2). ‘Chardonnay’ had more clusters, on average, in 1997 (84.1) and 1999 (79.5) than other years.

**Figure 1.**
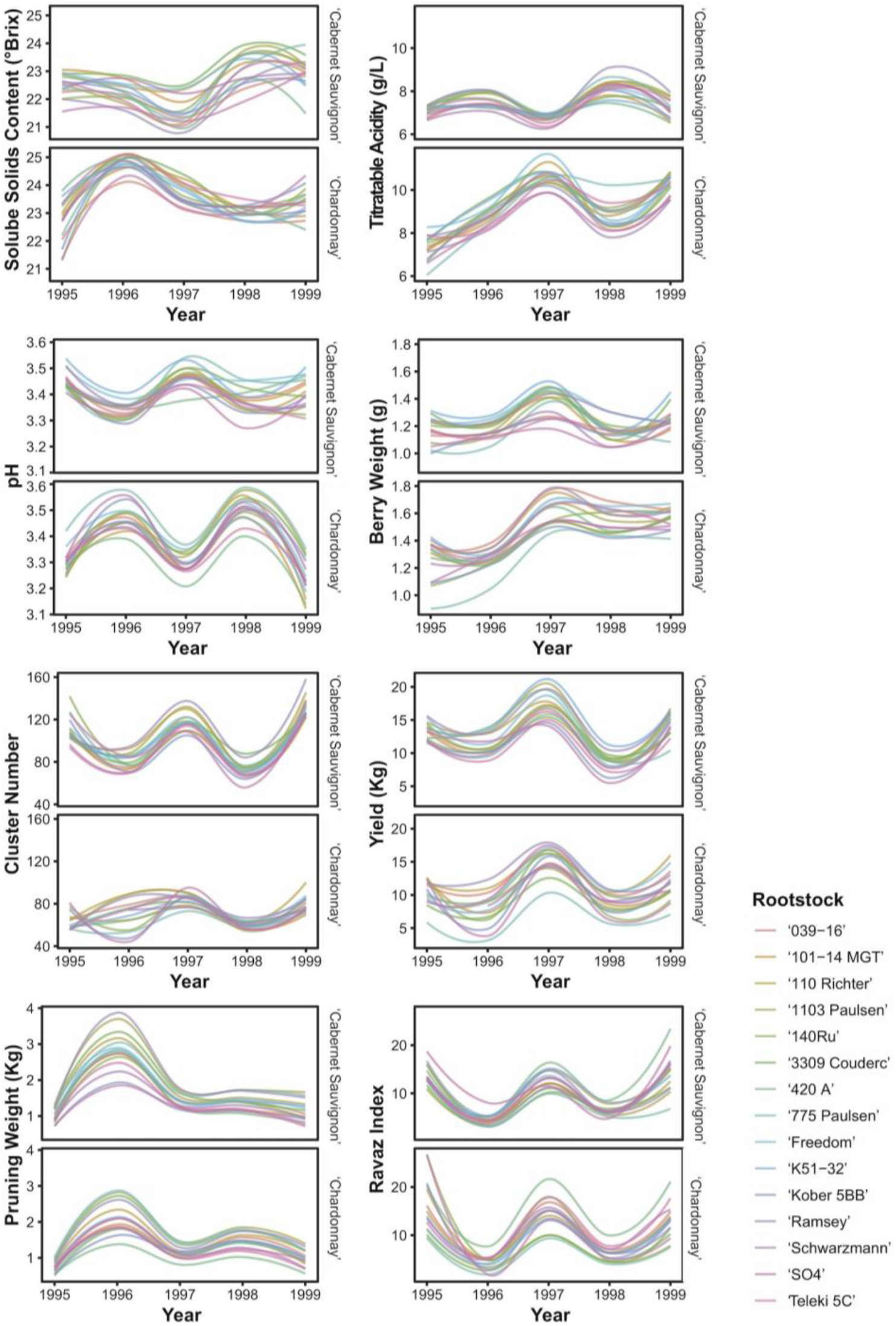
Phenotypic variation across years (1995 to 1999) for each rootstock by scion combination. Ravaz index is a measurement of crop load calculated by dividing yield by pruning weight from the following dormant season. Loess smoothing lines are plotted, however, the data are independent and these are for visualization purposes only. Individual data points for this figure are plotted in Figure S5.

In addition to decreased yields in 1996, vines also generally had higher pruning weights, with average values of 2.82 Kg for ‘Cabernet Sauvignon’ and 2.13 Kg for ‘Chardonnay’, in comparison to values ranging from 1.01 Kg to 1.43 Kg for ‘Cabernet Sauvignon’ and 0.74 Kg to 1.49 Kg for ‘Chardonnay’ in other years. However, the relative rankings of the rootstocks were generally consistent across years (Figure 1, Figure S5).

### Statistical modeling

Using a linear model (Equation 1), we identified year as the largest source of variation captured by our data (Figure 2). Year was a significant term for all phenotypes, explaining 8.46% (pH) to 52.04% (pruning weight) of the phenotypic variation. Year explained over 40% of the variation in pruning weight (52.04%), Ravaz index (48.64%), and yield (41.72%).

**Figure 2.**
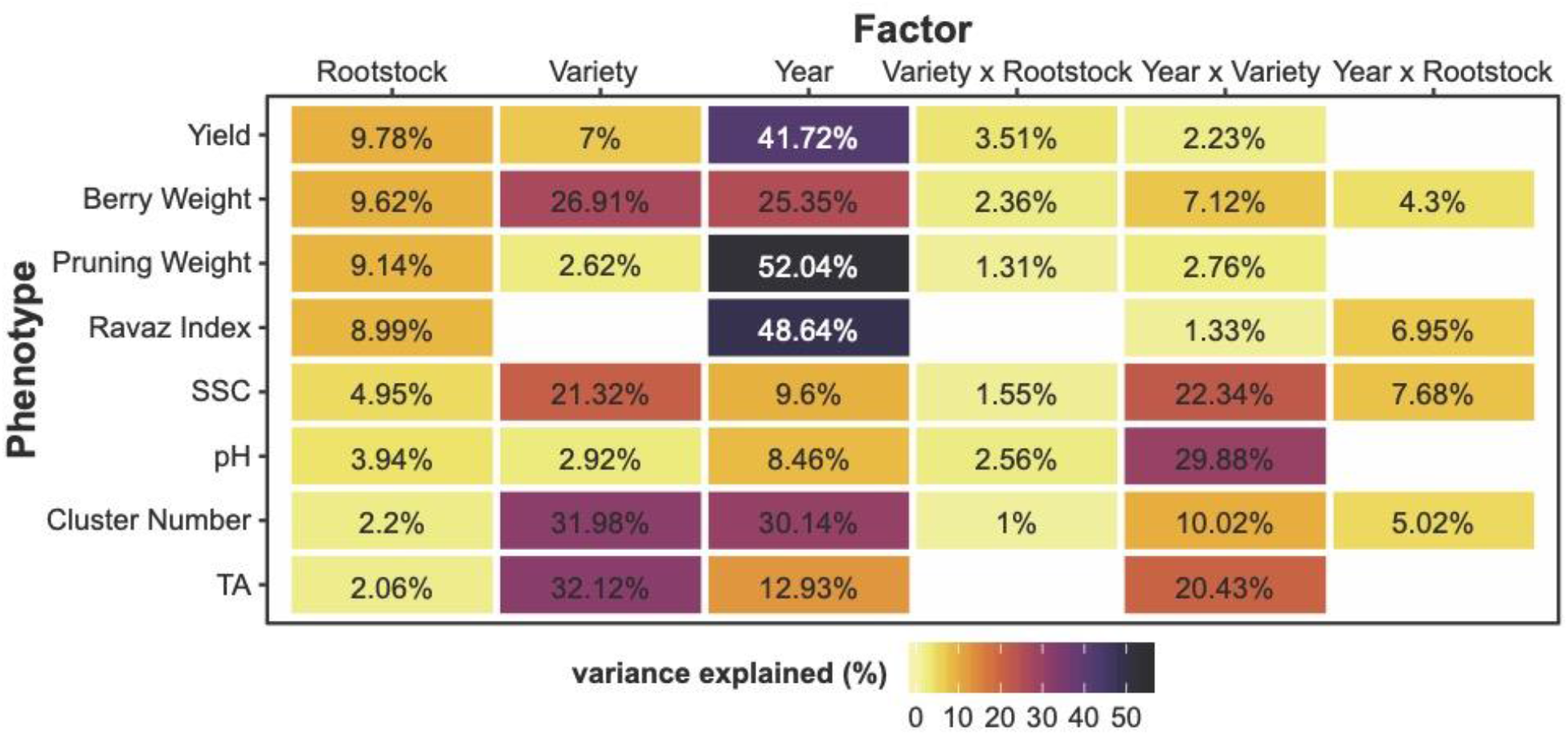
Phenotypic variation explained by factors of interest estimated using a linear model (equation 1). For each phenotype, the linear model was optimized by removing non-significant interaction effects. For factors which explain a significant amount of variance (p < 0.05), the percent variance explained is indicated using colour. Position in the vineyard (block) was included in the model but is not plotted. Phenotypes are sorted in order of the most variance explained by rootstock.

Grapevine variety explained a significant amount of variation in all traits except Ravaz index, with the strongest effect for TA (32.12%), cluster number (31.98%), berry weight (26.91%), and SSC (21.32%). The interaction between year and variety was significant for all traits, and over 20% of the variation in pH (29.88%), SSC (22.34%), and TA (20.43%) could be explained by this term.

Rootstock had a significant effect on all phenotypes and explained between 8.99% and 9.78% of the variation in yield, berry weight, pruning weight, and Ravaz index. For yield, pruning weight, pH, and TA, the interaction between rootstock and year was removed from the model because it did not explain a significant amount of variation in the trait. For the remaining traits, the interaction between rootstock and year explained 4.31% to 7.68% of the variation (Figure 2).

While the interaction between variety and rootstock was retained as a significant term for all phenotypes except TA and Ravaz index, it explained less than 4% of the variation in any given phenotype.

### Comparing rootstock performance

Focusing on the phenotypes in which rootstock showed the strongest effect, we plotted the distributions for yield (9.78%), berry weight (9.62%), pruning weight (9.14%), and Ravaz index (8.99%) and compared each of the 15 rootstocks using a Tukey test (Figure 3). Across these phenotypes, ‘Ramsey’ had among the highest yields, berry weights, and pruning weights, and one of the lowest Ravaz indexes. The yield for ‘Ramsey’ was significantly higher than eight of the other rootstocks evaluated. Similarly, ‘Freedom’ ranked within the top four for yield, berry weight, and pruning weight measurements, but ranked 11th for Ravaz index. However, ‘Freedom’ and ‘Ramsey’ only had a significantly lower Ravaz index than ‘420 A’.

**Figure 3.**
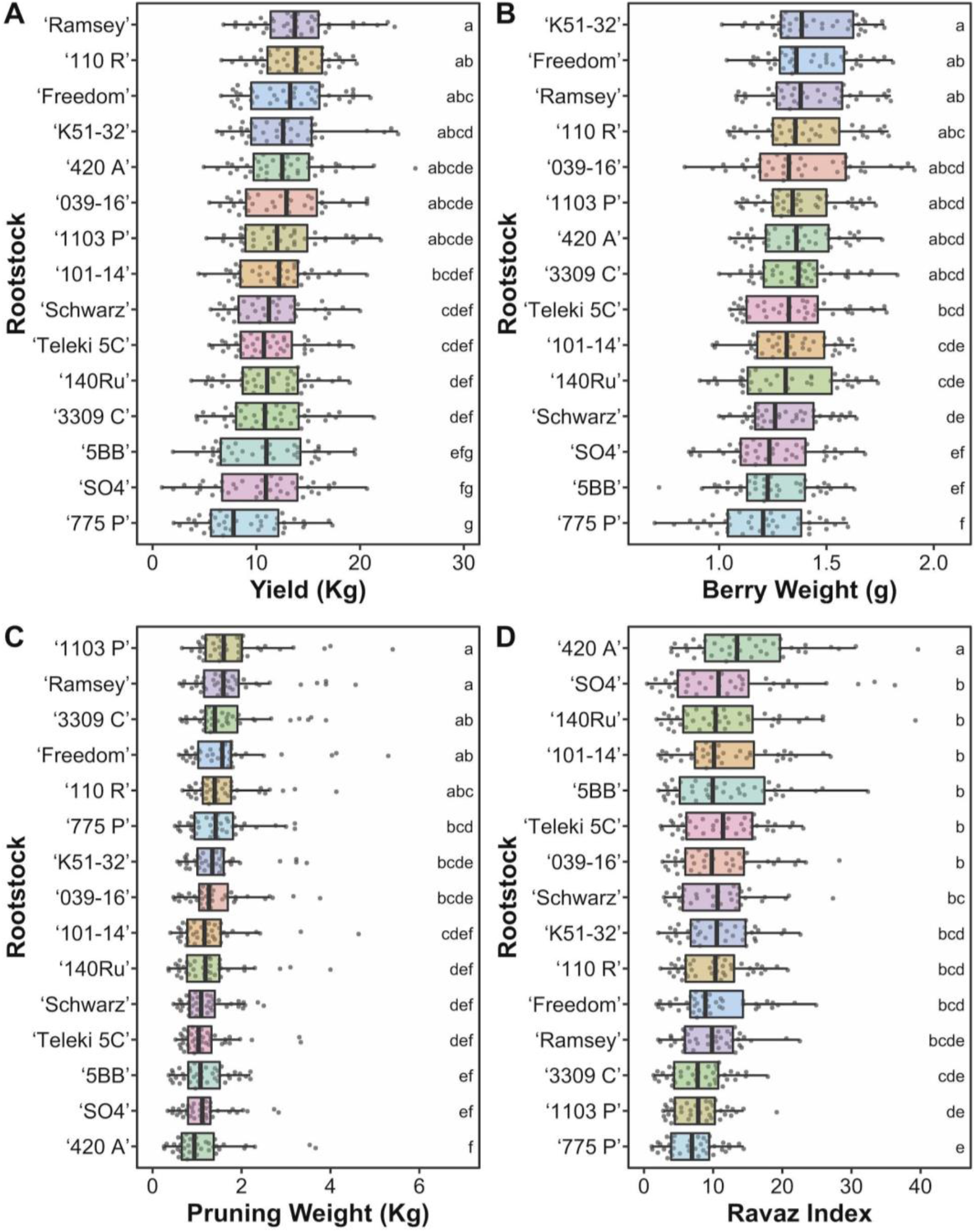
Variation in (A) yield, (B) berry weight, (C) pruning weight, and (D) Ravaz index across vines grafted to 15 different rootstocks. Rootstocks are ordered from highest to lowest mean values. Tukey test results are reported from a linear model accounting for variation in variety, year, position in the vineyard (block), and applicable interaction effects. Rootstocks with the same letter (indicated inside the plot) are not significantly different from each other. For estimated marginal means see Figure S3.

The rootstock ‘775 P’ generated the lowest yields and smallest berries, with only ‘SO4’ and ‘5BB’ not differing significantly for these two phenotypes. In contrast, ‘775 P’ was ranked 6th for pruning weight, which resulted in a significantly lower Ravaz index than all other rootstocks except ‘3309 C’, ‘1103 P’, and ‘Freedom’, although this trend is likely due primarily to the low yield of ‘Chardonnay’ grafted to ‘775 P’ (Figure S4). In comparison, ‘420 A’ ranked 5th for yield and had the lowest pruning weight, thus resulting in a Ravaz index which was significantly higher than all other rootstocks.

### Correlation between traits

For both ‘Chardonnay’ and ‘Cabernet Sauvignon’, Ravaz index was significantly correlated with most other phenotypes with the exception of TA (‘Chardonnay’) and SSC (‘Cabernet Sauvignon’) (Figure 4, Table S4). Ravaz index was positively correlated with cluster number for ‘Chardonnay’ (r = 0.250, p = 3.398 × 10^−4^) and ‘Cabernet Sauvignon’ (r = 0.671, p < 1 × 10^−15^). Yield was not significantly correlated with pruning weight for either variety but it was positively correlated with cluster number (‘Chardonnay’: r = 0.634, p < 1 × 10^−15^; ‘Cabernet Sauvignon’: r = 0.722, p < 1 × 10^−15^) and berry weight (‘Chardonnay’: r = 0.489, p < 1 × 10^−15^; ‘Cabernet Sauvignon’: r = 0.579, p < 1 × 10^−15^).

**Figure 4.**
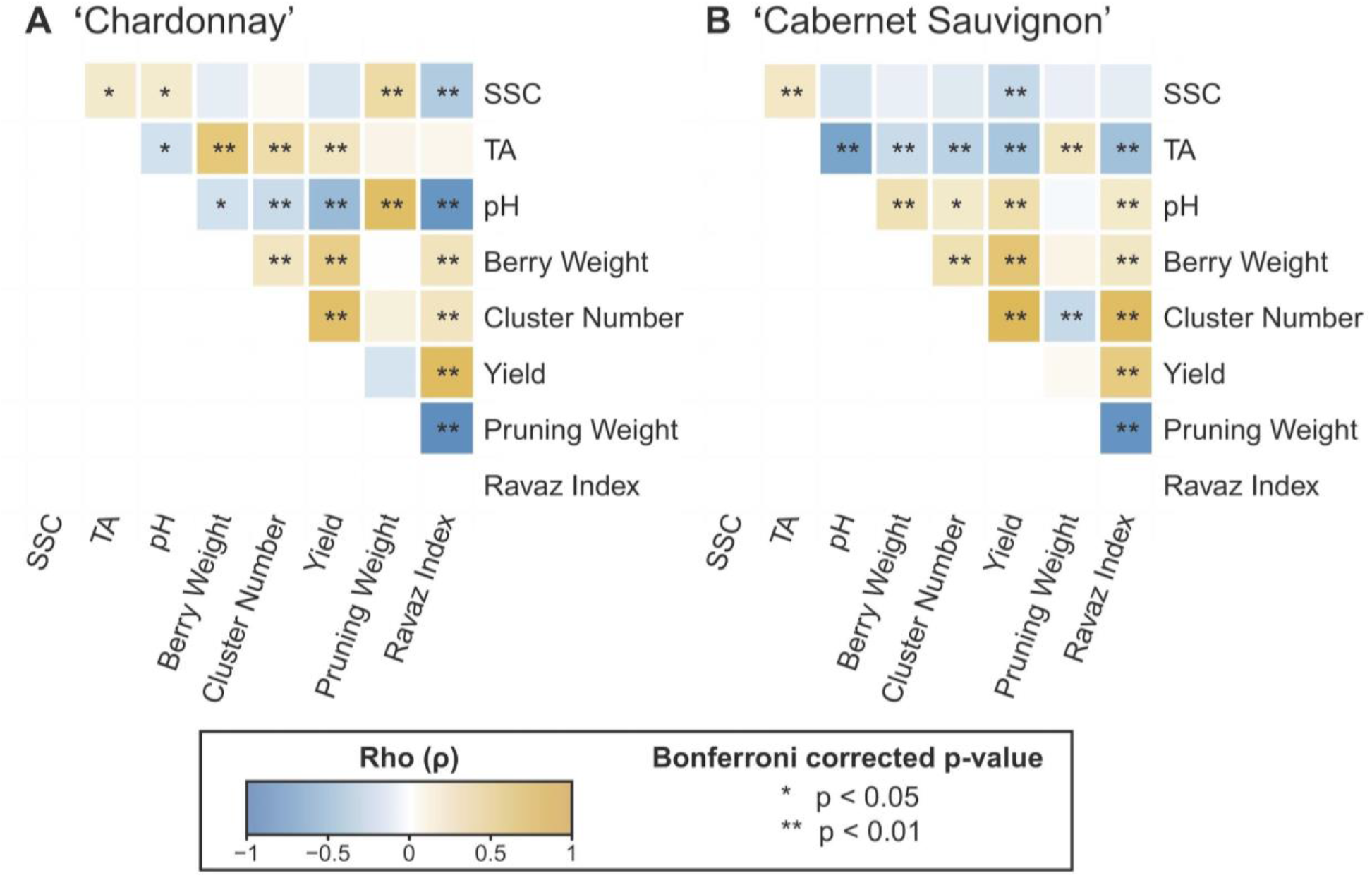
Spearman’s correlations among phenotypes for (A) ‘Chardonnay’ and (B) ‘Cabernet Sauvignon’. P-values were Bonferroni-corrected for multiple comparisons with a particular variety.

### Range of rootstock effects

Lastly, we evaluated the percent change between the best and worst performing rootstocks for each phenotype (Figure 5, Table S5). The percent change ranged from 3.10% for SSC to 93.78% for Ravaz index. In addition to Ravaz index, cluster number (35.42%), pruning weight (70.82%), and yield (77.35%) all increased by over 30%, while the remaining phenotypes increased by less than 15%.

**Figure 5.**
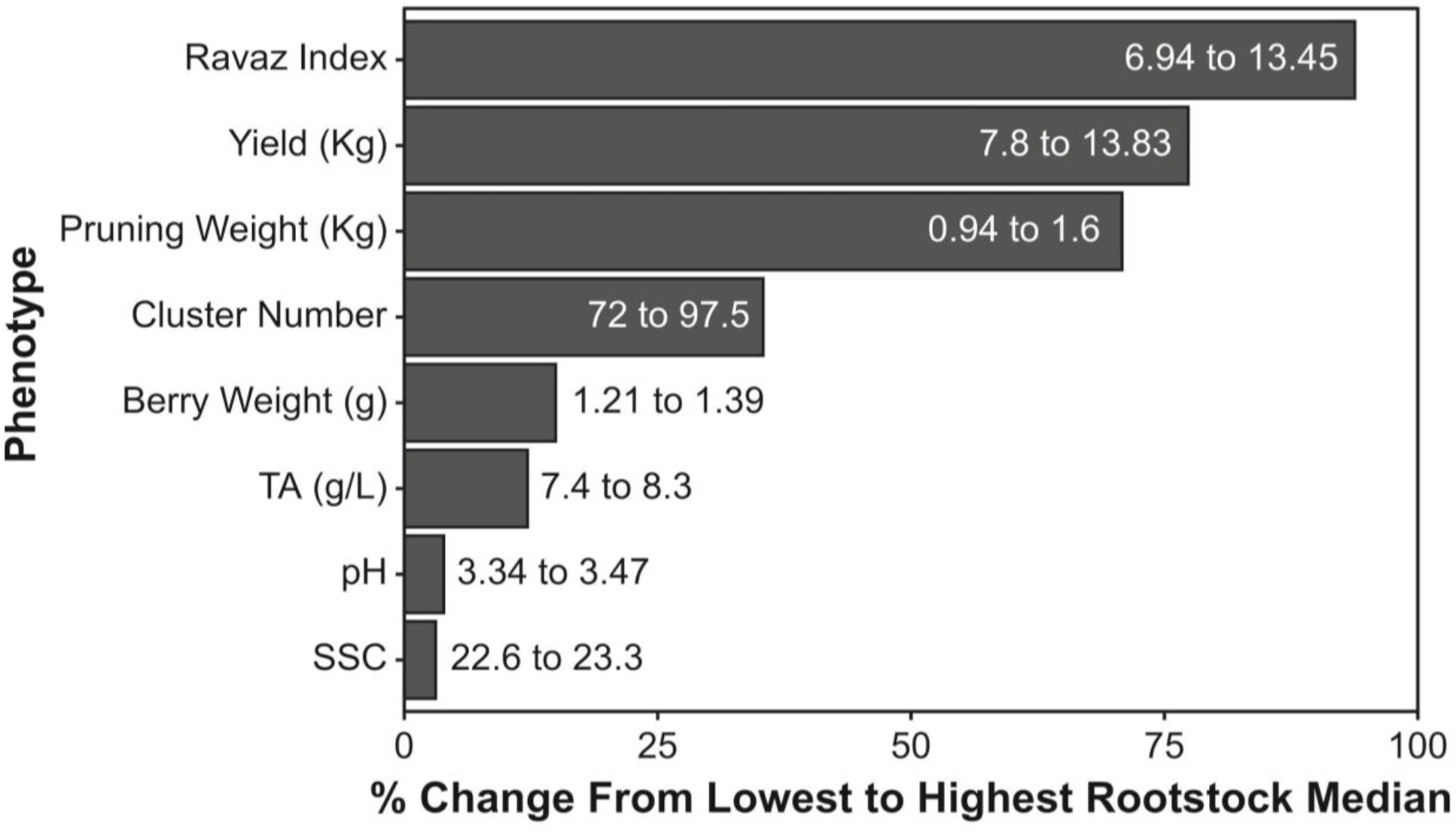
Percent change in each phenotype from rootstock with the lowest median to the rootstock with the highest median. Phenotypes are ordered from largest percent change to lowest percent change. Raw values are also listed.

## Discussion

### Potential causes of variation across years

In California, most wine regions annually receive sufficient rainfall during the dormant season to support desired canopy growth. However, there are years where low dormant season rainfall may reduce canopy growth^31^. Previous work examining ‘Merlot’ vines across two years in California found that soil moisture level during the dormant season impacted both vegetative and reproductive growth, even when irrigation is applied after budbreak^31^. In our study, there was less rainfall during the 1994 and 1997 dormant seasons when floral initiation would have occurred (Figure S2). While the reduction in yield we observed in 1998 may be due to a dry 1997 dormant season, the dormant season prior to 1996 had higher rainfall and it’s unclear why the yield was not higher (Figure 1). Thus, at least for the 1998 growing season, a reduction in rainfall the previous year may have had a more severe impact on reproductive growth in contrast to vegetative growth.

In addition to the impact of rainfall, overcropping (too much fruit) has been considered as a source of alternate (biennial) bearing, with high yields in one year reducing yield in the subsequent year. Growers may be concerned that mechanical pruning can lead to overcropping, however, the vines in this study were hand-pruned to a target of 50 nodes per vine. In addition, previous work on both ‘Sultana’ and ‘Concord’ grapes found variation in yield was primarily due to environmental factors and management practices, rather than alternate bearing because of overcropping^32,33^. Therefore, in our study, it was most likely the low precipitation and not overcropping in 1997 which was primarily responsible for reducing yields in 1998.

In contrast to our study (Figure 1), previous work found that the reduction to pruning mass due to dormant season rainfall was more severe than the reduction to yield, increasing the Ravaz index for vines which did not receive rainfall^31^. While this is not consistent with our results, we do find that year is the largest source of variation in growth-related traits (Figure 2), confirming that variable environmental conditions between years, such as access to water during the dormant season, plays a crucial role in plant growth and development.

### Variation and consistency of scion and rootstock effects

During the growing season, grape growers can use management practices such as irrigation to partly buffer against year-to-year variation^34^. The vines in this study were all irrigated using the same management practices across five years, with a moderate RDI program of 80% from berry set to veraison, therefore reducing the impact of weather fluctuations during the growing season. When included in a linear model (Figure 2), variety explained over 20% of the variation we observed for TA, berry weight, and SSC, indicating that there is a strong variety-specific effect on many berry characteristics. Year, or vintage, had a significant interaction with variety for all phenotypes and explained over 20% of the variation in berry chemistry measurements such as pH, SSC, and TA. Even with consistent water management, berries from each variety responded differently to environmental conditions. In comparison, for growth measurements such as yield and pruning weight there was less variation explained by year by variety interaction, indicating the years with low or high growth for ‘Chardonnay’ had a similar impact on ‘Cabernet Sauvignon’. Thus, the effect of year on growth was relatively similar across different grapevine varieties, while the effect on berry chemistry differed between varieties.

In contrast to variety, the effect of rootstock rarely varied across years (Figure 2): the interaction between rootstock and year was not a significant term for yield, pruning weight, pH or TA, and explained less than 8% of the variation for the remaining phenotypes. We found that the effect of a rootstock was generally consistent between ‘Chardonnay’ and ‘Cabernet Sauvignon’, with the interaction between variety and rootstock explaining very little phenotypic variation (less than 4%). This suggests that grape growers should place great emphasis on rootstock choice as a critical decision during vineyard planning as performance of one rootstock, relative to others, is generally consistent over time and between varieties.

### Effects of rootstocks and their interaction with environment

The choice of rootstock is particularly important for growth-related traits such as yield, pruning weight, berry size, and Ravaz index, where rootstock effects explained at least 9% of the variation (Figure 2). In contrast to our study, previous work examining nine grape varieties grown ungrafted and grafted to four different rootstocks found that yield and berry weight were not affected by rootstock^35^. However, similar to our work, the study identified that vine and yield components were more responsive to rootstock than fruit composition variables^35^. Our results are also consistent with previous work identifying a significant difference in yield, pruning weight, and berry weight of ‘Shiraz’ vines grafted to different rootstocks and measured across six years^18^. That said, while rootstock can have a significant impact on yield, environmental factors including location, climate, and soil, generally have a much larger influence on this trait^19,36^.

In long-lived perennial plants where significant year-to-year variation can occur, the collection of data across multiple years is a valuable tool for untangling the effect of the environment. By evaluating the vines in this study across five years, we were able to account for the variation due to year in our model and determine how much of the variation was due to rootstock (Figure 3). Similarly, a recent seven year study examined ‘Cabernet Sauvignon’ grafted to three different rootstocks. The study found no significant effect of rootstock on pruning weight, although yield and berry weight did differ significantly^37^. When comparing the rootstocks which overlapped with our study, the authors found similar results: ‘101-14 Mgt’ and ‘420 A’ did not differ significantly for yield and berry weight, but ‘420 A’ had a significantly higher Ravaz index^37^.

A 25 year study that measured ‘Cabernet Sauvignon’ grafted to three different rootstocks found that Ravaz index was significantly affected by rootstock choice, but only after 7 years of planting. Similarly, yields across rootstocks only differed after 15 years^38^. Although we detect variation in vines which had been planted for three to seven years, our dataset includes a much broader representation of rootstocks. The effects observed in our study may not only be due to the young age of the vines examined here, but also due to the particular soil of the experimental plot. For example, previous work found that in a vineyard severely infected with nematodes, grafting ‘Chardonnay’ to 15 different rootstocks increased yield by up to 7 times and pruning weight by up to 23-fold when compared to ungrafted vines. Rootstocks varied in their resistance and tolerance to nematodes, with rootstock parentage influencing both yield and pruning weight^21^. Given that grapevines may remain in the ground for at least 20 years, additional long term studies across multiple locations are needed in order to determine how the effect of rootstocks changes over time and under different external conditions including environment and soil.

### Rootstocks affect growth-related scion phenotypes

Generally, rootstocks resulting in large values of one growth-related phenotype also resulted in large measures of other growth-related phenotypes (Figure 4). For example, rootstocks that generated higher yields generally also produced larger berries. While cluster number was more highly correlated with yield than berry weight, much more variation in berry weight could be explained by rootstock, indicating that increased yields due to rootstock were primarily a result of increased berry weight and not additional clusters. This suggests that rootstock choice does not influence floral initiation, but rather influences water uptake, which leads to variation in berry weight. While high yields are generally desirable, the ratio of skin-to-pulp is an important consideration for vinification, and this ratio is reduced when berries take on more water. Previous work also demonstrated that in addition to decreasing fruit size, reducing water in ‘Cabernet Sauvignon’ increased desirable characteristics such as the concentrations of skin tannin and anthocyanins^39^. Therefore, while the use of a rootstock to increase yields is beneficial, this has to be balanced with ensuring that the berries maintain a desirable size, possibly through a reduction in irrigation for more vigorous rootstocks.

While increased reproductive growth leading to increased yields is economically beneficial, if the vegetative growth increases at the same rate, the Ravaz index, or crop load, of the vine will remain consistent. Increased vegetative growth results in higher vine management costs, such as pruning and leaf thinning. We demonstrated that Ravaz index was correlated with most of the other phenotypes we measured (Figure 4). This suggests that the balance between reproductive and vegetative growth in a vine is associated with many other characteristics of that vine. However, pruning weight and yield were not correlated, likely because all vines were pruned to a similar size and shoot number to prevent overcropping, but the number of clusters per shoot differed. As a result, higher yields were positively correlated with both berry weight and cluster number, but the correlation with cluster number was higher for both varieties, indicating that the primary source of increased yield was more clusters and not larger berries. In some instances, therefore, rootstock choice may increase reproductive growth of a vine without an increase in vegetative growth and its associated costs.

The choice of one roostock over another can result in nearly a two-fold difference in growth-related traits like yield, Ravaz index and pruning weight but has little effect on berry on the chemistry measurements assessed in this study, in particular SSC and pH (Figure 5). Previous studies have also found small differences in berry chemistry such as SSC with large variation in growth, such as yield, due to rootstock21,43.

### Potential causes of rootstock-induced variation in growth

While we were unable to evaluate it directly, we find it likely that much of the variation in growth that can be attributed to rootstock in our study is due to increased water uptake by vines grafted to certain rootstocks. Variation in water uptake is generally the result of some combination of water uptake efficiency, the size and surface area of the root system, and stomatal regulation to reduce water loss, among other factors^12^. For example, ‘Ramsey’ and ‘Freedom’ generally had high yields, large berries, and high pruning weights (Figure 3). Similarly, in an Australia study, ‘Shiraz’ vines grown with irrigation and grafted to ‘Ramsey’ or ‘Freedom’ rootstocks yielded more fruit than ungrafted vines and than vines grafted on the other five rootstock varieties assessed, indicating that these rootstocks tend to increase yield and pruning weight^40^. Other work found a rootstock-dependent effect of irrigation on some yield components such as cluster number and berry weight, but not on yield itself ^41^. In our study, all vines were irrigated equally, which may have led to a rootstock-specific effect on water uptake which ultimately contributed to variation in yield and could be further controlled with rootstock-specific irrigation regimes.

In addition to variation water uptake, it is possible some variation in growth is due to variation in disease resistance. While phylloxera is a concern in the region, all vines were grafted to rootstocks which should provide protection. Additionally, there is the potential for grapevine fanleaf virus at this site. One of the key symptoms of fanleaf degeneration is a decrease in fruit set which leads to a lower yield^42^. Given that only vines grafted to ‘039-16’ would have fanleaf protection in this study, and the yield of vines grafted to ‘039-16’ is not significantly higher than other rootstocks (Figure 3A) which do not offer protection, indicating that it is likely not a severe concern in this vineyard. Thus, while there may be some variation in rootstock tolerance to other pests and pathogens, this is unlikely to be a major factor in this study.

### Conclusion

Increasing yield, especially during the early years of production, can have a dramatic influence on the profitability of a vineyard and the results of this study clearly indicate that selection of the right rootstock is a valuable tool that grape growers can use to help control vine size and yield. These results should be taken into account when considering which rootstock to select, particularly in the San Joaquin Valley where this study was performed, with additional work needed to verify the effect of each rootstock across different geographic locations. Future work can explore if the early advantage provided by rootstock is maintained throughout the life of a vineyard.

## Supplemental information

**Figure S1. Vineyard map of rootstock evaluation trial**.

**Figure S2. Variation in maximum temperature (°F), minimum temperature (°F) and cumulative precipitation (inches) measured from January 1994 to December 1999 in Lodi, California, US**.

**Figure S3. Estimated marginal means with 95% confidence interval for variation in (A) yield, (B) berry weight, (C) pruning weight, and (D) Ravaz index across vines grafted to 15 different rootstocks. Rootstocks are ordered from highest to lowest mean values. Linear models Tukey test results are reported from a linear model accounting for variation in variety, year, position in the vineyard (block), and applicable interaction effects**.

**Figure S4. Median values (+/- standard deviation) for each phenotype for ‘Chardonnay’ and ‘Cabernet Sauvignon’ grafted to each rootstock**.

**Figure S5. Phenotypic variation across years (1995 to 1999) for each rootstock by scion combination with individual data points plotted. Loess smoothing lines are also plotted, however, the data are independent and these are for visualization purposes only. Ravaz index is a measurement of crop load calculated by dividing yield by pruning weight from the following dormant season**.

**Table S1. Harvest and pruning dates for ‘Chardonnay’ and ‘Cabernet Sauvignon’ vines sampled from 1995-1999**.

**Table S2. Phenotype data collected from 1995 to 1999 for ‘Chardonnay’ and ‘Cabernet Sauvignon’ vines grafted to 15 different rootstocks**.

**Table S3. Linear model results for each phenotype**. Each model was optimized for each phenotype: the main effects were retained in all cases but non-significant interactions were removed.

**Table S4. Results of Spearman’s correlation between phenotypes for ‘Chardonnay’ and ‘Cabernet Sauvignon’**. P-values were Bonferroni-corrected for comparison within each variety.

**Table S5. Variation across phenotypes based on median rootstock values**. The maximum median, minimum median, average median are included as well as the maximum percent change (from minimum to maximum median) and average percent change across rootstocks.

## Data Availability

All data supporting the results of this manuscript have been included as supplementary materials (Table S2). The code used to analyze the data and generate the figures in this manuscript has been made publicly available on GitHub^27^.

## Acknowledgments

ZM was supported by the National Science Foundation Plant Genome Research Program 1546869. SM was supported by the National Sciences and Engineering Council of Canada. We thank Matthew Rubin (Donald Danforth Plant Science Center) for statistical expertise. We gratefully acknowledge all of the individuals involved in maintaining the vineyard evaluated for this study, in particular Ernie and Jeff Dosio (Pacific AgriLands Inc., Modesto, CA).

## Authors’ Contributions

PV conceived this work and led the collection of the data. LMJ organized the data. ZM analyzed the data and wrote the manuscript. PC, SM, and DHC provided feedback in data analysis and interpreting the results. PC, RKS, and DHC provided project oversight and guidance. All authors reviewed the manuscript.

