## Supplementary material for "Grapevine rootstocks affect growth-related scion phenotypes": Figure S1

### Experimental Vineyard

Spacing: 7x10 ft 2.13x.3.05 m

| **Block** | 9 vines | 8 vines | 8 vines | 8 vines | ↑N |
|---|---|---|---|---|---|
| **Variety** | **Rootstock** | | | | **Row** |
| Cabernet Sauvignon | K51-32 | | Schwarzmann | | 30 |
| | 101-14 MGT | Schwarzmann | 110 Richter | K51-32 | 29 |
| | 1103 Paulsen | 420 A | Ramsey | Teleki 5C | 28 |
| | 140 Ruggeri | 775 Paulsen | K51-32 | 039-16 | 27 |
| | 420 A | 3309 Couderc | Freedom | Freedom | 26 |
| | 110 Richter | Kober 5BB/S04 | 101-14 MGT | 420 A | 25 |
| | 3309 Couderc | K51-32 | 039-16 | Kober 5BB/S04 | 24 |
| | Kober 5BB/S04 | 101-14 MGT | 775 Paulsen | 1103 Paulsen | 23 |
| | Schwarzmann | 140 Ruggeri | Teleki 5C | 140 Ruggeri | 22 |
| | K51-32 | 1103 Paulsen | 3309 Couderc | 110 Richter | 21 |
| | Teleki 5C | 039-16 | Kober 5BB/S04 | Schwarzmann | 20 |
| | 775 Paulsen | Ramsey | 1103 Paulsen | 101-14 MGT | 19 |
| | 039-16 | Freedom | Schwarzmann | 3309 Couderc | 18 |
| | Ramsey | Teleki 5C | 140 Ruggeri | Ramsey | 17 |
| | Freedom | 110 Richter | 420 A | 775 Paulsen | 16 |
| Chardonnay | 101-14 MGT | 3309 Couderc | 110 Richter | 039-16 | 15 |
| | Schwarzmann | 140 Ruggeri | Teleki 5C | K51-32 | 14 |
| | Kober 5BB/S04 | Ramsey | 101-14 MGT | Freedom | 13 |
| | 110 Richter | Freedom | 039-16 | 420 A | 12 |
| | 3309 Couderc | Schwarzmann | K51-32 | Kober 5BB/S04 | 11 |
| | Ramsey | 1103 Paulsen | 3309 Couderc | 1103 Paulsen | 10 |
| | K51-32 | Teleki 5C | 420 A | 140 Ruggeri | 9 |
| | Freedom | 039-16 | 775 Paulsen | 110 Richter | 8 |
| | 775 Paulsen | Kober 5BB/S04 | Ramsey | Schwarzmann | 7 |
| | Teleki 5C | 420 A | Schwarzmann | 101-14 MGT | 6 |
| | 039-16 | K51-32 | 1103 Paulsen | 3309 Couderc | 5 |
| | 1103 Paulsen | 110 Richter | 140 Ruggeri | Ramsey | 4 |
| | 140 Ruggeri | 101-14 MGT | Freedom | 775 Paulsen | 3 |
| | 420 A | 775 Paulsen | Kober 5BB/S04 | Teleki 5C | 2 |
| | K51-32 | | Schwarzmann | | 1 |

Parking Area/ Fence
