## Supplementary figures and images for "Grapevine rootstocks affect growth-related scion phenotypes"

### Figure S2

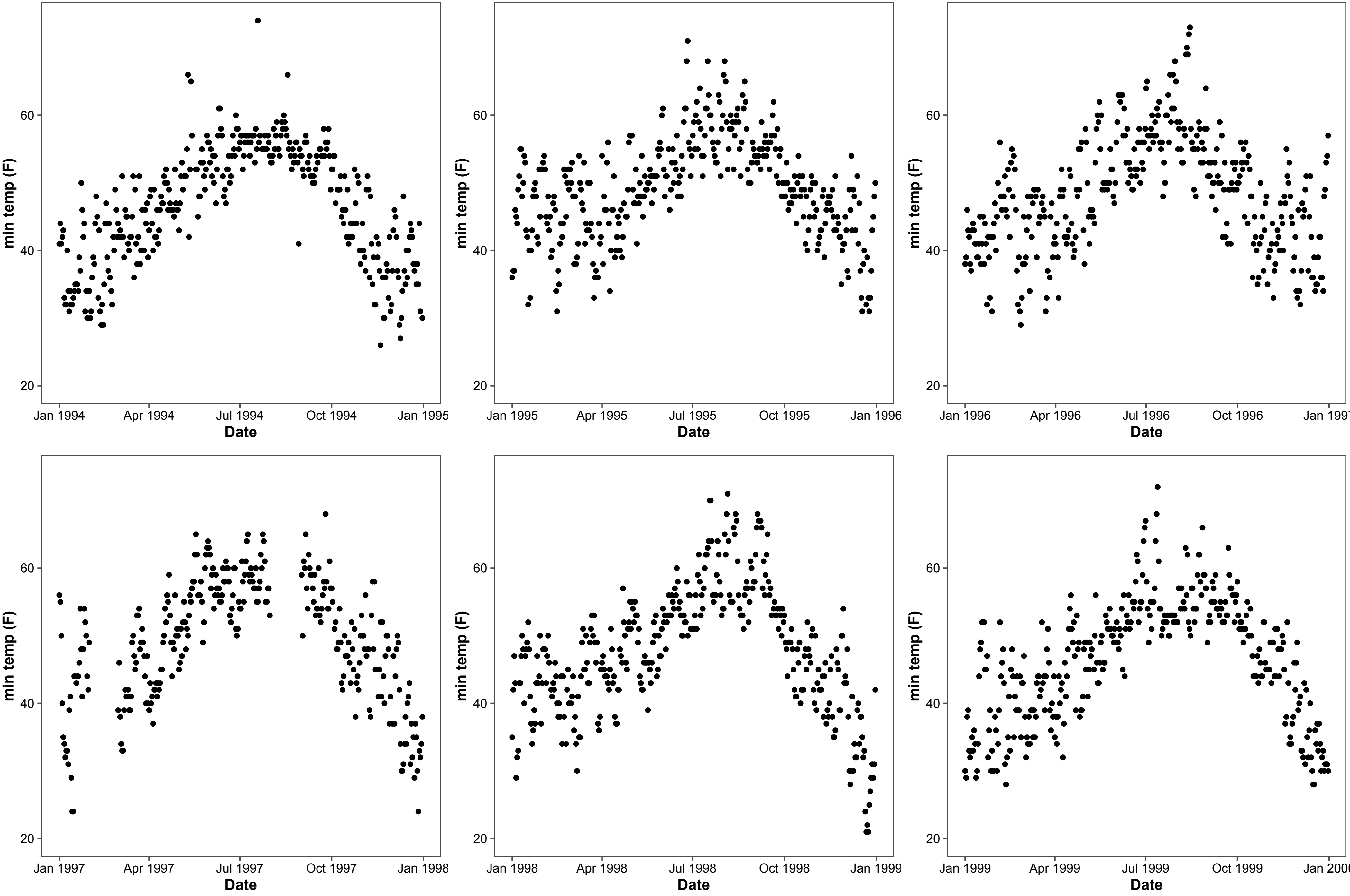

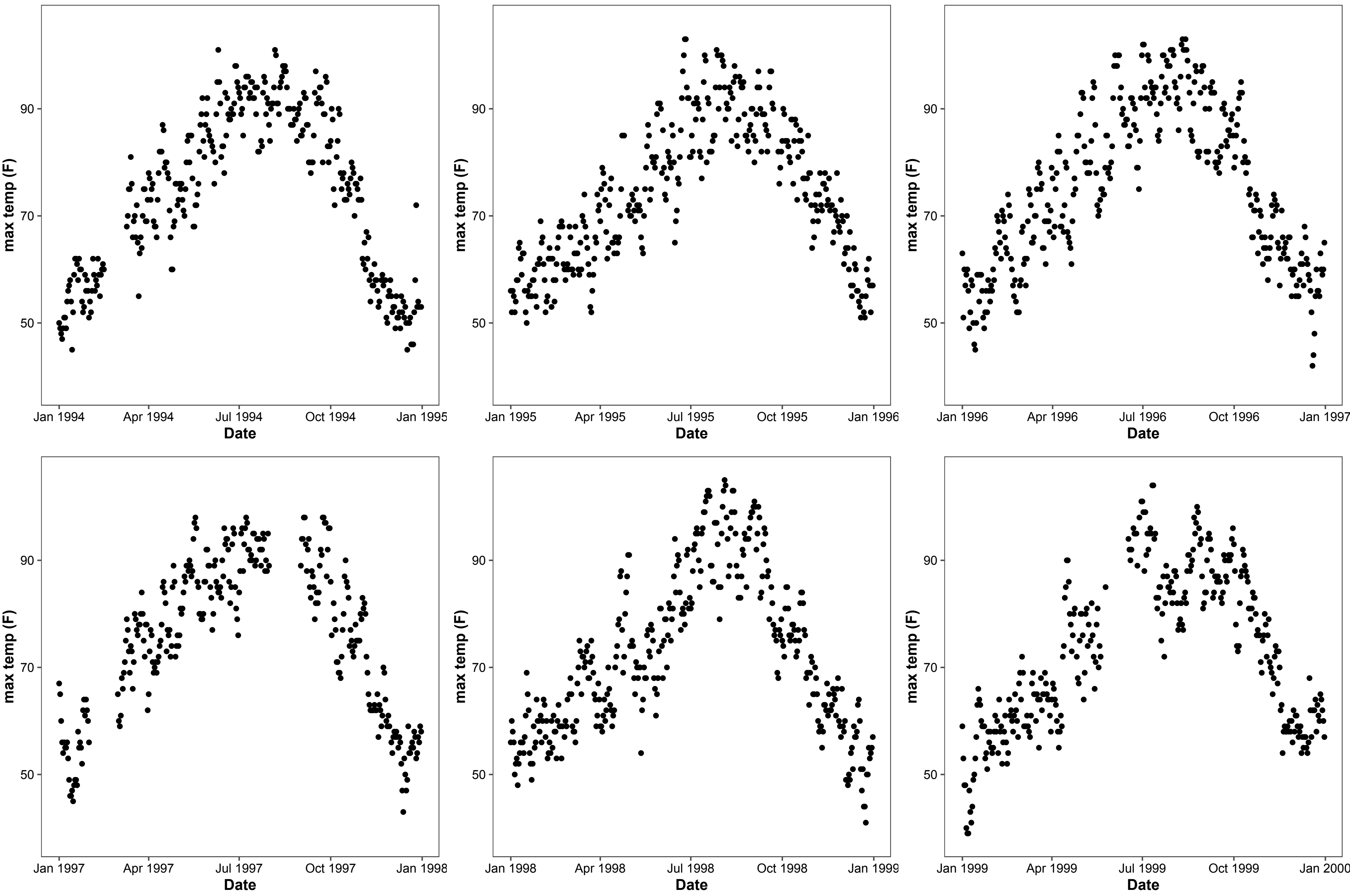

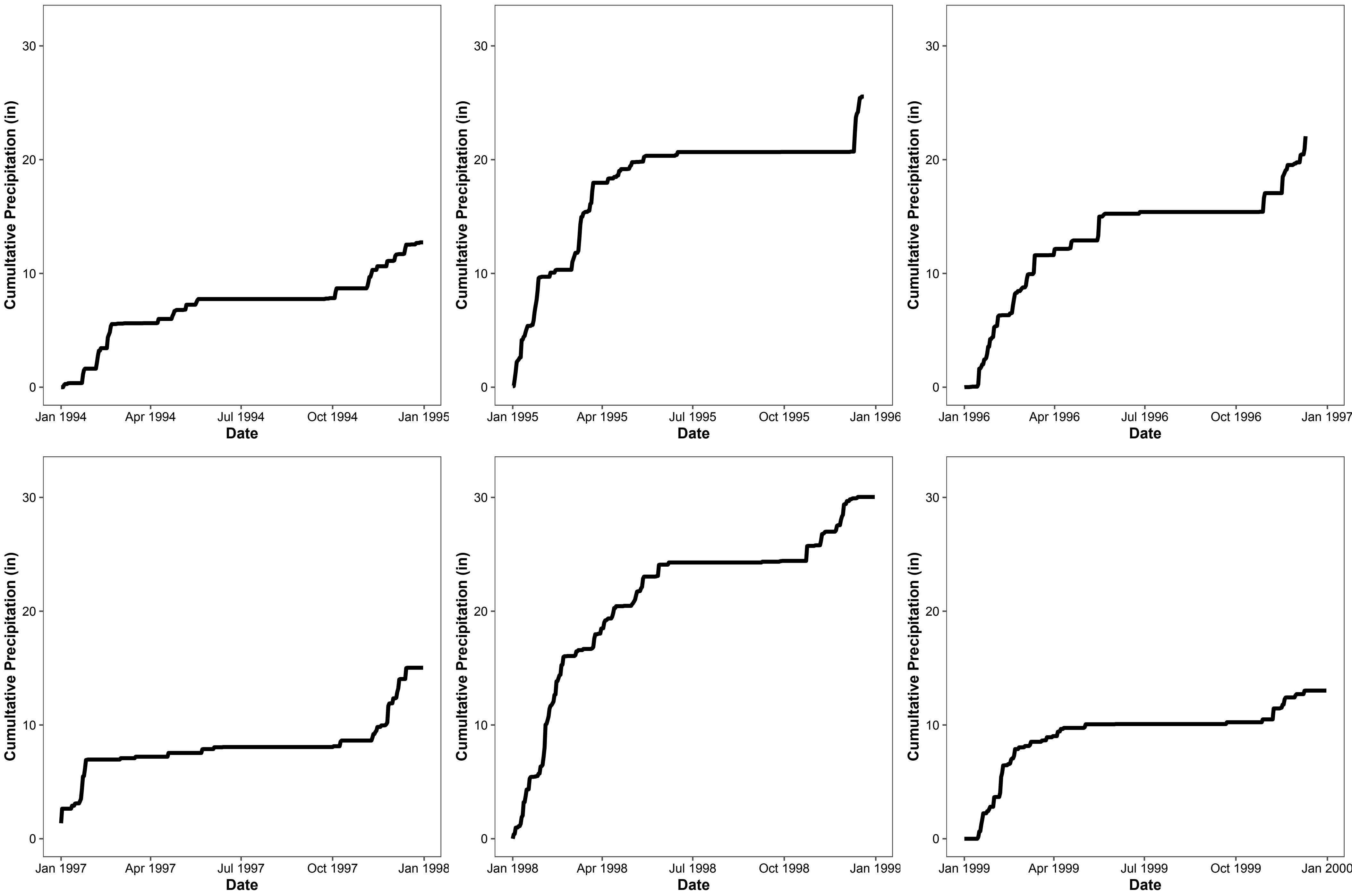

### Figure S3

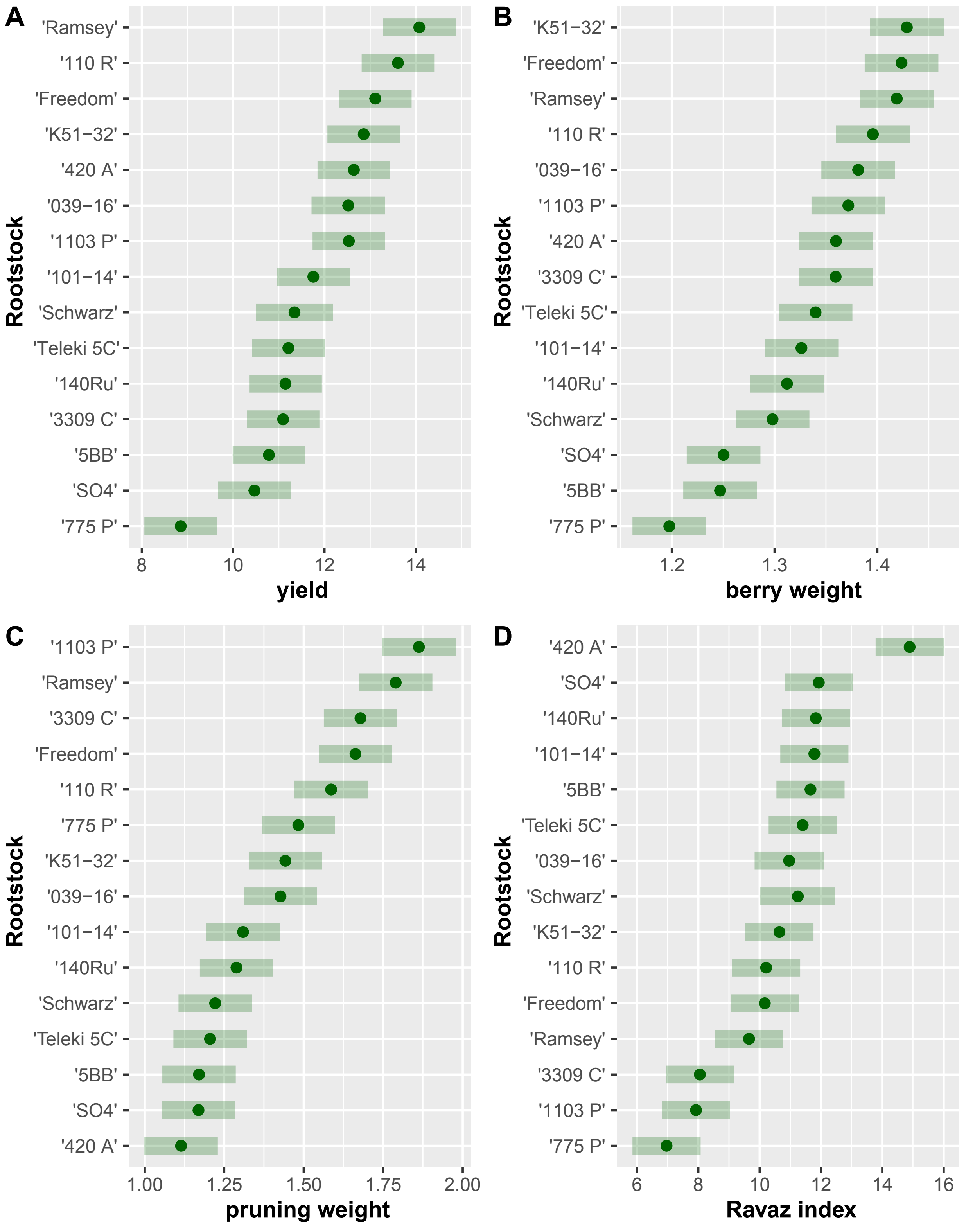


**Estimated marginal means with 95% confidence interval**

### Figure S4

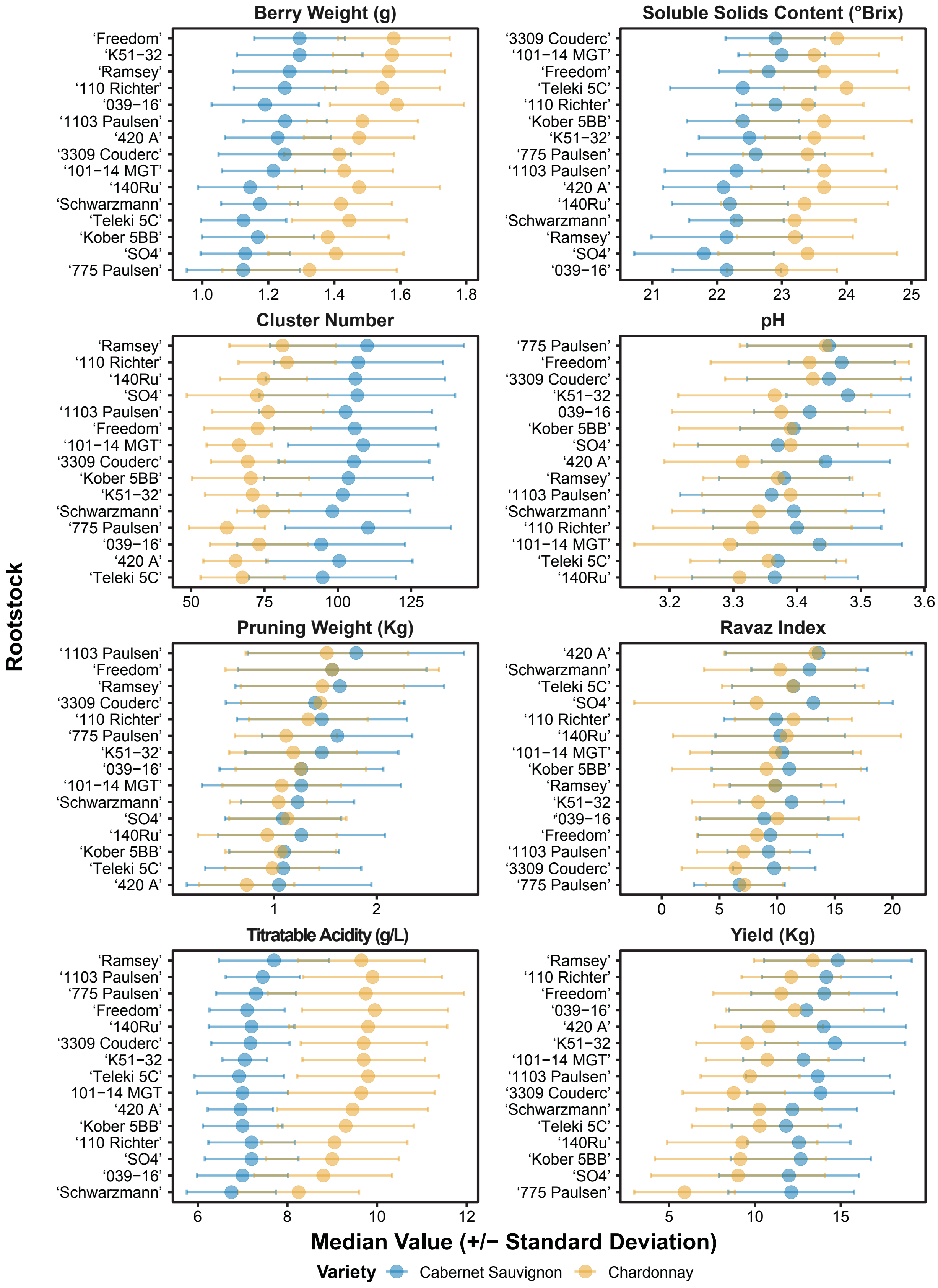

### Figure S5

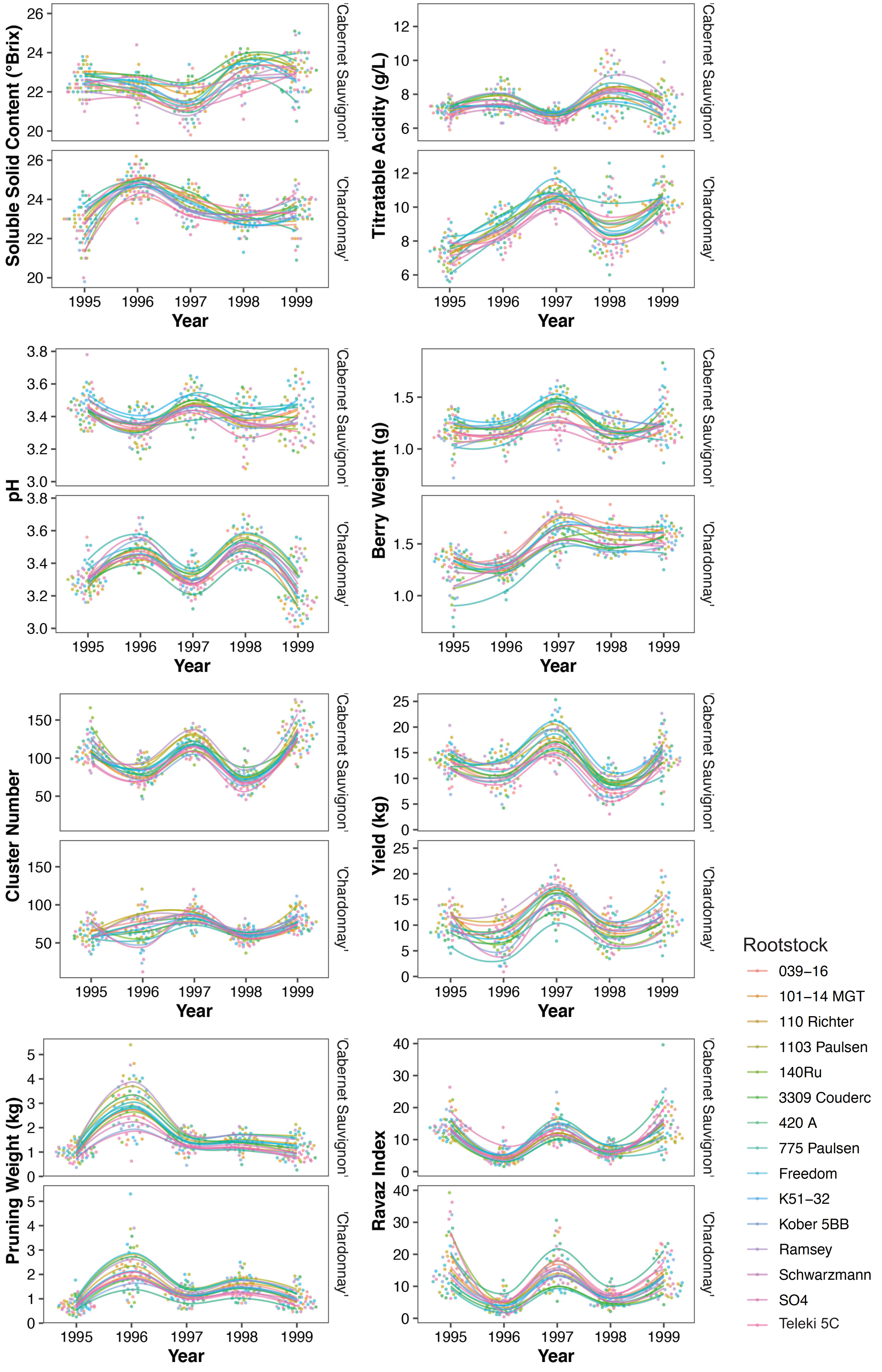
